# Rasal1 impairment unleashes anticancer immunity - a focus on T cells

**DOI:** 10.64898/2026.01.19.700452

**Authors:** Mark E. Issa, Alejandro Schcolnik-Cabrera

## Abstract

Immune checkpoint blockade (ICB) has demonstrated clinical efficacy in several cancers, including melanoma, lung, colorectal, and liver malignancies. However, a substantial proportion of patients fail to respond, underscoring the need for alternative immunotherapeutic strategies capable of overcoming resistance to conventional checkpoint inhibition. One such strategy involves targeting intracellular inhibitory immune checkpoints that regulate effector lymphocyte function. Rasal1, a Ras GTPase-activating protein, has been shown to negatively regulate T cell-mediated antitumor immunity. In this study, we further characterized the impact of Rasal1 impairment on tumor progression, T cell stemness, and effector function. Using an endonuclease-mediated mutation targeting the C2 domain of Rasal1, we demonstrate that Rasal1-impaired (Rasal1i) mice exhibit significantly reduced tumor growth across multiple murine cancer models. Rasal1i mice displayed increased intratumoral CD8^+^ T cell accumulation, activation, cytolytic capacity, and enhanced Wnt signaling. Tumor-infiltrating lymphocytes additionally exhibited increased progenitor and stem-like memory phenotypes. Notably, Rasal1 inhibition prolonged survival and potentiated αPD-1 therapy in a resistant PD-L1-expressing B16F10 melanoma model. Collectively, these findings identify Rasal1 as an intracellular inhibitory immune checkpoint that constrains T cell stemness and antitumor function, and support its further evaluation as a therapeutic target for cancer immunotherapy.

## 1. INTRODUCTION

Immune checkpoint blockade (ICB) has significantly improved the effectiveness of cancer treatments. However, the inhibition of PDL1, PD1 or CLTA4 has demonstrated success with only a proportion of cancer patients. This is possibly due to the development of resistance, whereby cancer cells alter PDL1 expression and render PDL1 inhibition non-effective. This limitation therefore necessitates the identification of an alternative immunotherapeutic strategy effective in non-responsive cancer patients. One such strategy could be the inhibition of intracellular checkpoints that negatively regulate immune cell responses[1-3].

Upon tumor antigen recognition by the T cell receptor (TCR), and the recognition of MHC-I by the CD8 coreceptor (or MHC-II by the CD4 coreceptor), p56 Lck is recruited to CD8 whereby it phosphorylates the tyrosine residues on the ζ chain’s ITAM, thereby creating docking sites for ZAP70[4]. ZAP70 phosphorylates the adaptor complex LAT, which creates docking sites for the adaptors SLP76, ADAP-SKAP1 complex, and the cell growth factor PLCγ1[5]. PLCγ1 catalyzes the production of the second messengers IP3 and DAG, thereby resulting in the mobilization of calcium signaling, the activation of the guanine exchange factor (GEF) RasGRP1, as well as the activation of NFκB and NFAT pathways[6]. Moreover, LAT recruits the adaptor protein GRB2 along with the GEF SOS to activate the small GTPase p21 Ras. RasGRP1 and SOS cooperate to stimulate a GTP-bound p21 Ras. Altogether, the MAPK pathway communicates cell growth signals from the cytoplasm to the nucleus in response to TCR activation[7]. In contrast to GEFs, GTPase-activating proteins (GAPs) stimulate a GDP-bound p21 Ras, thereby rendering it inactivated. The Ras activator-like 1 (Rasal1) protein belongs to the Rasa GAP subfamily which includes NF1, Rasa1-4 and Rasal1-3, SynGAP and DAB2ip[8]. Rasal1 is composed of two C2 domains, followed by a GAP catalytic domain, a pleckstrin homology (Ph) domain and a Bruton tyrosine kinase (Btk) motif. Rasal1 is directed to the plasma membrane in response to intracellular calcium increase through its C2 domains, which anchor Rasal1 to membrane phosphatidylinositol lipids, an event that triggers GAP activity and inhibition of Ras[7,9].

In T cells, Rasal1 exerts its effects specifically by favoring the inactive GDP-bound p21 Ras form and thereby inhibiting the MAPK pathway[7]. Rasal1 has been shown to negatively regulate T cell antitumor immunity in which Rasal1 knockdown results in a significant decrease in B16F10 melanoma pulmonary metastases, and a significant decrease in EL4 lymphoma growth. These effects were accompanied by an increase in CD8^+^ tumor infiltrating lymphocytes (TILs), an increase in the CD8^+^ T cell granzyme B, and an increase in CD8^+^ T cell interferon-γ (IFN-γ)[7].

In the present study, we investigated the effects of Rasal1 impairment on anticancer immunity, while focusing on T cell stemness in murine cancers. Using an endonuclease-mediated Rasal1 impairing mutation, termed Rasal1i (aka Rasal mutant), we demonstrate that Rasal1i reduced skin cancer growth in both males and females, colorectal cancer in females and lung cancer in males; prolonged survival and potentiated αPD1 in resistant PDL1-expressing B16F10 tumor model. Rasal1 mutant mice showed augmented CD8^+^ T cell functions in B16F10 melanoma. Rasal1i also resulted in an increase in B16F10 intratumoral CD8^+^ T cell presence, T cell activation, antigen experience, TCR-Ras signaling in T cells, and CD8^+^ T cell production of cytolytic enzyme granzyme B. Furthermore, Rasal1i promoted Wnt signaling in CD8^+^ T cells in B16F10 melanoma and resulted in a significant increase in stem-like memory T cells in cancer. Overall, this study delineates the role of Rasal1 as a negative intracellular immune checkpoint, and as a key player in the regulation of T cell progenitor phenotype.

## 2. MATERIALS AND METHODS

### 2.1 Animal housing and ethical approval

All experiments were approved by the CR-HMR, aka Comité de protection des animaux du CIUSSS de l’Est-de-l’Île-de-Montréal (F06 CPA-21061 du projet 2017-1346, 2017-JA-001-2). All mice employed for this project were housed at the Hôpital Maisonneuve Rosemont animal facility (Montréal, QC, Canada). Up to 5 mice per cage were housed in ventilated cages under a 12h light/dark cycle, with cleaning of cages twice per week, and access to food and water *ad libitum*.

### 2.2 Mice

C57BL/6N mice were purchased from Jackson Laboratory. The Rasal1 impaired (Rasal1i) mouse model was purchased from the mouse biology program in California (https://www.mmrrc.org/). The Rasal1i allele is an endonuclease-mediated impairment intragenic mutation. The Rasal1i allele was generated at the Jackson Laboratory by injecting Cas9 RNA a n d t h e f o u r g u i d e s e q u e n c e s A G G C T A G A G T C T C A T A T G G T, T G G G G G C AT G T G TA G A C T C T, G A G A AT G G G C T G AT T C A C A G a n d TGAACCGAATCACATATCAA, which resulted in a 481 bp deletion spanning exons 4-5 beginning at chromosome 5 positive strand position 120, 654, 693 bp, TCTAGGAGTGTGTGTGTGAC, and ending after GAGGGAGAATGGGCTGATTC at 120, 655, 173 bp (genome of reference: GRCm38/mm10). This mutation deletes exons 4 and 5, and 305 bp of flanking intronic sequence including the splice acceptors and donors. This deletion is predicted to cause a change of amino acid sequence after residue 41 and early truncation 14 amino acids later. Offspring were genotyped as per the provider’s instructions (MMRRC:049371-UCD) on a regular basis using the protocol provided by the Jackson Laboratory. *For additional information on the phenotype of the Rasal1i allele, readers are invited to follow (https://www.informatics.jax.org/allele/MGI:5774569, https://www.mousephenotype.org/data/genes/MGI:1330842, https://www.mmrrc.org/catalog/sds.php?mmrrc_id=49371)*.

### 2.3 Cell culture

Splenocytes-derived T cells were cultured in RPMI-1640, whereas B16F10, B16F10-PDL1, MC38, and LLC1 cells were cultured in DMEM. All medium used was supplemented with 10% FBS (Gibco^TM^, MT35077CV) and 1% penicillin/streptomycin (Gibco^TM^, 15140122). B16F10 cells were split every 2 or 3 days, and were maintained in a standard humidified atmosphere supplemented with 5% CO_2_ at 37°C. The cell lines derive from ATCC, except for the B16F10-PDL1 cell which was kindly donated to us by Dr Michel Ardolino of the University of Ottawa, ON, Canada[10]. Cells were routinely tested for *Mycoplasma*.

### 2.4 Experimental design in vivo

Male and female mice, aged 8-12 weeks, were implanted with tumor cells after a 7 day-period of acclimation. For the MC38 and LLC1 cancers, males and females were implanted with 500,000 cells subcutaneously in 100 µL serum-free medium in the right flank. The B16F10 or the B16F10-PDL1 melanoma cell line were intradermally inoculated using 27 ½ gauge syringes, unless otherwise indicated. The number of cells inoculated in males was 100,000 cells in 100 µL serum-free medium in the right flank, and in females it was 50,000 cells in 50 µL serum-free medium in the right flank, following [11]. Tumor measurements were initiated at day 4 post implantation using a caliper, and measures were taken every second day until the end of the experiment. At the end of the experiment, mice were anesthetized with isoflurane and then euthanized by cervical dislocation. Tissues of interest were then harvested. In accordance with IACUC protocols, Human endpoints were implemented to minimize animal suffering if the following signs of distress were found: diminished movement, pale palms, or hinged backs.

### 2.5 T cell stimulation and qPCR

Spleens were resected from healthy mice, and tumors were collected diseased mice, placed immediately on ice. Tissues were crushed on 70 µM cell strainer, and single cell suspensions were collected in serum-free RPMI-1640 medium. Cells were centrifuged at 300*g* for pelleting. Splenocytes were depleted from red blood cells (RBCs) as per manufacturer’s instructions (*the full list of antibodies and kits is shown in **S. Table 1***). RBC-depleted splenocytes were plated and stimulated with αCD3ε/αCD28 at 2 µg/mL each, incubated for an indicated period, and used for downstream application. T cell stimulation was verified by the appearance of T cell clusters in the plates and by cell numbers using trypan blue. RT-qPCR methods were performed according to the TRIzol RNA extraction procedures suggested detailed by the provider Invitrogen. RNA was quantified using NanoDrop, and subsequently, RT and qPCR were performed according to Abm standard protocols (https://www.abmgood.com/all-in-one-5x-rt-mastermix.html, https://www.abmgood.com/blastaq-2x-qpcr-mastermix.html) on an AB Biosystems StepOne real time PCR system. The full list of primer sequences is shown in **S. Table 2**.

### 2.6 CFSE proliferation assay in vitro

CFSE proliferation assays were conducted as per manufacturer’s instructions. Briefly, RBC-depleted splenocytes were incubated in a 1 µM working CFSE solution at 10^7^ cells/mL for 20 minutes at room temperature protected from light. Staining was quenched adding 5 times the original staining volume of cell culture medium containing 10% FBS. CFSE-labeled cells were pelleted and resuspended at 2 x 10^6^ cells/mL (2mL/well in 24 well plates). Splenocytes were stimulated with αCD3ε/αCD28 at 2 µg/mL each, incubated for 48 hours, and thereafter proliferation was assessed by flow cytometry.

### 2.7 Tumor-infiltrating lymphocyte collection

Tumors were harvested and minced into small pieces before enzymatic digestion with Liberase TL and DNase1 at 100 µg/mL each, at 37^°^C for 30 min. Addition of RPMI-1640 with 10% FBS stopped the reaction, and the resulting mixture was passed through 70 µm cell strainers to generate single cell suspension. After 2 washes with 1X PBS, cells were fractionated into (i) total TILs for quantification proportion from whole tumor homogenates, or (ii) overlayed on a ficoll layer for lymphocyte collection. Total tumoral tissues and ficoll-concentrated lymphocytes were then stained for viability and fixed with 2% PFA according to manufacturer’s protocol. Whole tumor homogenates and enriched TILs were then stored at -80 °C until further processing.

### 2.8 Flow cytometry

Flow cytometric analyses were performed according to standard protocols. Standard tested antibodies with verified reactivity were purchased from BioLegend or eBioscience. In brief, PFA-fixed TILs were washed twice with ice-cold 1X PBS and stained with 1 µL of manufacturer’s working antibody concentration for surface targets for 30 min on ice. For intracellular staining, cells were permeabilized for 45 min on ice, and staining were performed using the FoxP3/Transcription Factor Staining Buffer Set (eBiosciences, San Diego, CA, USA). TILs were stained with 2 µL of manufacturer’s working antibody concentration for intracellular targets for 45 min on ice, and then washed twice with PBS and proceeded to data collection. Cells were acquired using BD LSR Fortessa X-20 and the DIVA software. FACS analyses were performed using FlowJo software version 10 and Cytobank premium for expression levels and colocalization of targets of interest.

### 2.9 Gating strategies

Subsets of immune cells were gated as shown in **S. Table 3**.

### 2.10 Bioinformatics analysis and patient stratification

Transcriptomic (RNA-seq) and matched clinical data from TCGA-SKCM (melanoma), TCGA-COAD/READ (colorectal adenocarcinoma), and TCGA-LUAD/LUSC (non-small cell lung cancer) were retrieved using TCGAbiolinks (v2.30+). STAR-aligned counts were converted to FPKM, log₂-transformed with a +1 pseudocount, and RASAL1 expression (ENSG00000111344) extracted from the first matching row. Sample types were classified using barcode positions 14–15 (01 = primary tumor, 11 = solid tissue normal). For survival analysis, only cases with valid data (non-missing time-to-event, time >0 days) were included. Overall survival (OS) was calculated from diagnosis to death (event=1) or last follow-up (censored=0), converted to years (365.25). Patients were stratified into high/low RASAL1 groups by cohort-specific median. Kaplan–Meier curves were compared using the log-rank test; univariate Cox models estimated hazard ratios (HR) with 95% CI (low expression as reference). Differential expression between tumor and normal samples was assessed by Wilcoxon rank-sum test on log₂(FPKM+1) values. All analyses were performed in RStudio using packages *survival*, *survminer*, *dplyr*, and *ggpubr*.

### 2.11 Protein expression analysis

Representative immunohistochemistry images of RASAL1 protein expression in malignant tissues were retrieved from the Human Protein Atlas (www.proteinatlas.org; version 25). Images were selected to reflect the most common/average staining patterns reported for melanoma, colorectal adenocarcinoma, and lung adenocarcinoma in patient-derived punch biopsies.

### 2.12 Statistical analysis

For each *in vivo* experiment, the depicted results derive from at least four mice per treatment group, except for Rasal1i assays with or without αPD1 experiments whereby the presented results derive from three mice per group. Analyses were performed with GraphPad Prism software version 10 (GraphPad Software, CA, USA). All data are expressed as mean ± SD, except AUC were expressed as mean ± SEM. A two-tailed t-test was used when only two groups were compared, and a one-way ANOVA with Turkey correction was used when more than two groups were compared. The difference in mean values between groups was significant when *p* < 0.05 *; *p* < 0.01 ** and *p* < 0.001 ***.

## 3. RESULTS

### 3.1 Rasal1i augments T cell activation and proliferation

We first addressed the effect of Rasal1i on T cell activation and proliferation *in vitro*. Freshly collected WT and Rasal1 impaired (Rasal1i) splenocytes were stimulated for 48 hours with αCD3ε/αCD28 at 2 µg/mL each. Cell counts were performed and results showed that stimulated Rasal1i splenocytes displayed increased numbers as compared to WT (**S. Fig. 1A**). Using the CSFSE proliferation assay, results also showed that stimulated Rasal1i splenocytes augmented CD8^+^ T cell proliferation as compared to WT, as depicted in the additional peak of the CFSE diagram (**S. Fig. 1A**). Stimulated Rasal1i CD8^+^ T cell splenocytes displayed increased expression of the activation markers CD44 and CD69, and lower expression of the inhibitory markers CTLA4 and PD1, as well as elevated levels of inflammatory cytokines, when compared to WT splenocytes (**S. Fig. 1B-D**). By examining the Immunogen database (immugen.org), it was found that inflammatory cytokines IL1β, IL6 and TNFα alter the expression of Rasal1 in murine splenocytes, corroborating the notion that Rasal1 plays an immune check role (**S. Fig. 1E**). Altogether, these data show that Rasal1i increases T cell activation and proliferation *in vitro*.

### 3.2 Rasal1i reduces cancer growth in mice

To address the effects of Rasal1i on tumor growth, tumor growth was observed over time, and the area under the curve (AUC) was computed. It was found that Rasal1i mitigated tumor growth LLC1 lung cancer and MC38 colorectal cancers (**Fig. 1A-B**), as well as in B16F10 melanoma in both males and females (**Fig. 1C-D**). The results also demonstrate in a visual manner that Rasal1 impairment mitigate tumor growth as depicted in the tumor photos in mouse B16F10 melanoma in males (**Fig. 1E**). An indicator for the degree of response *in vivo* is depicted in a pie chart (**Fig. 1F**). Altogether, these data demonstrate that Rasal1i reduces murine cancer growth.

**Fig. 1.**
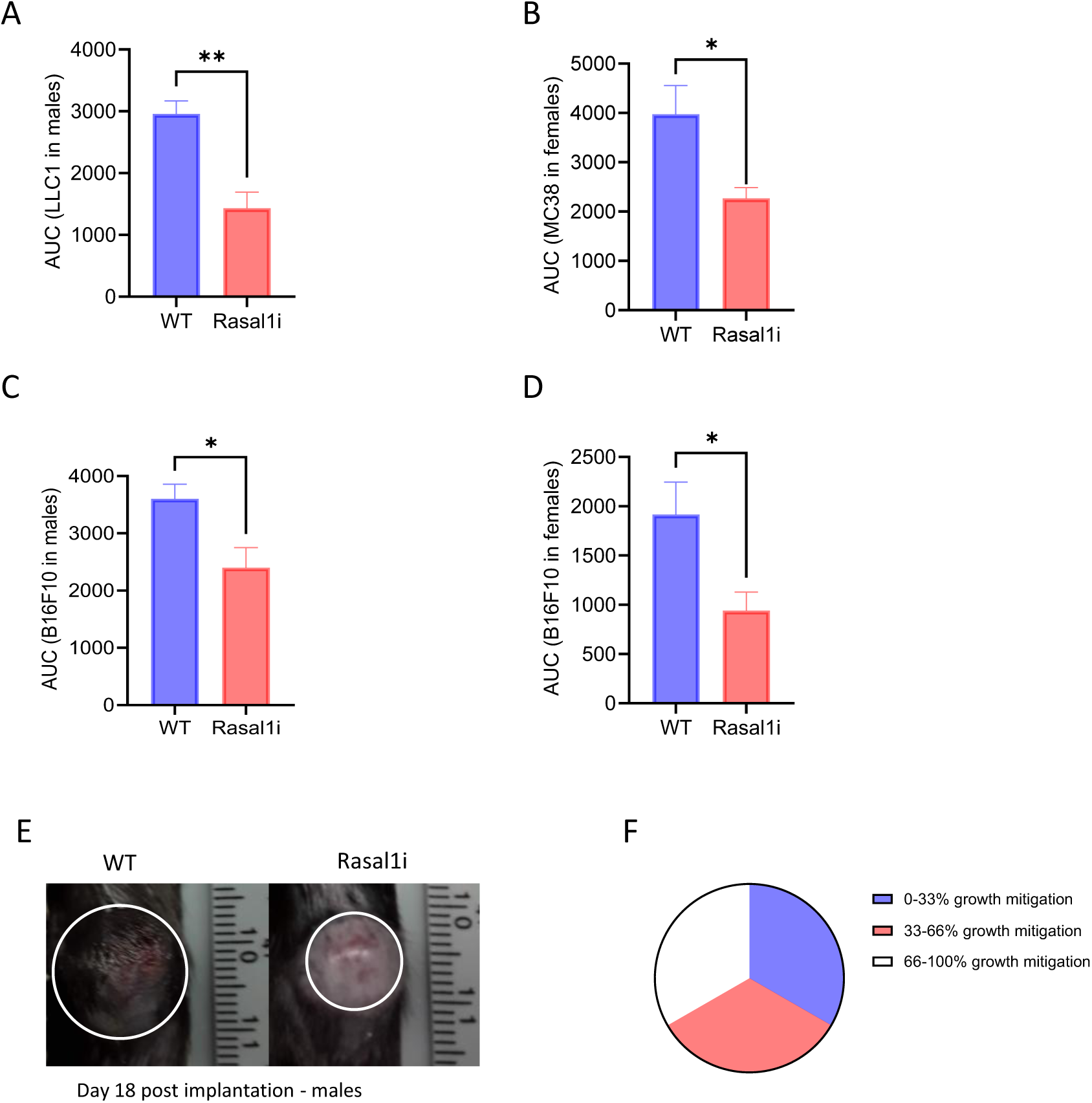
Rasal1i reduces tumor growth in mice. AUC for LLC1 lung cancer growth in males, and MC38 colorectal cancer growth in females (**A & B**), B16F10 skin cancer growth in males and females, respectively (**C & D**). Representative images of tumor growth of B16F10 melanoma in males (**E**). An approximation scale for *in vivo* degree of response (**F**). Data shows a representative experiment for each cancer model. *B16F10 was conducted four times in males, two times in females; B16F10-PDL1 one time in males and one time in females, LLC1 one time in males, and MC38 two times in females*. *For AUC statistical analyses, parametric unpaired two-tailed t-tests were used. *p < 0.05, **p < 0.01 and ***p < 0.001*.

### 3.2 Rasal1i results in augmented immune and T cell infiltration

Next, we isolated the TILs and examined their composition. For the majority of the study, and for immunological purposes whereby we wanted to an easily accessible orthotopic model, we chose to work mostly with the B16F10 model in male mice. B16F10 intratumoral immune cell presence was substantially increased in Rasal1i melanoma-bearing mice when compared to WT mice (**Fig. 2A**). This result was corroborated in LLC1 and MC38 tumors (**S. Fig. 2A-B**). The proportion of T cells within the overall immune cell compartment was increased in response to Rasal1i (**Fig. 2B**). Rasal1i resulted in a significant shift towards CD8^+^ T cells within the T cell compartment, as represented by an increase in the CD8^+^ TIL percentage with a concomitant decrease in CD4^+^ TILs within T cells in males (**Fig. 2C-D**). Rasal1i CD8^+^ TILs also showed an increased expression of the Ras pathway targets CD44 and Ki67 (**Fig. 2E-F**). Furthermore, a viSNE analysis corroborated the observation that the T cell compartment was shifted towards CD8^+^ T cells in response to Rasal1i (**Fig. 2G**). To help the reader visualize the potential biochemical interactions within CD8^+^ TILs, a representation of the interacting pathways relevant to this article are depicted in **Fig. 2H**. Furthermore, an interrogation of Rasal1 expression in other hematopoietic cells revealed that Rasal1 is expressed in several lineages, including but not limited to, T cells, B cells, and dendritic cells (DCs) (https://www.ebi.ac.uk/gxa/home) (**Fig. 2I**). Taken together, these results indicate that Rasal1 impairment results in an expanded intratumoral immune cell compartment.

**Fig. 2.**
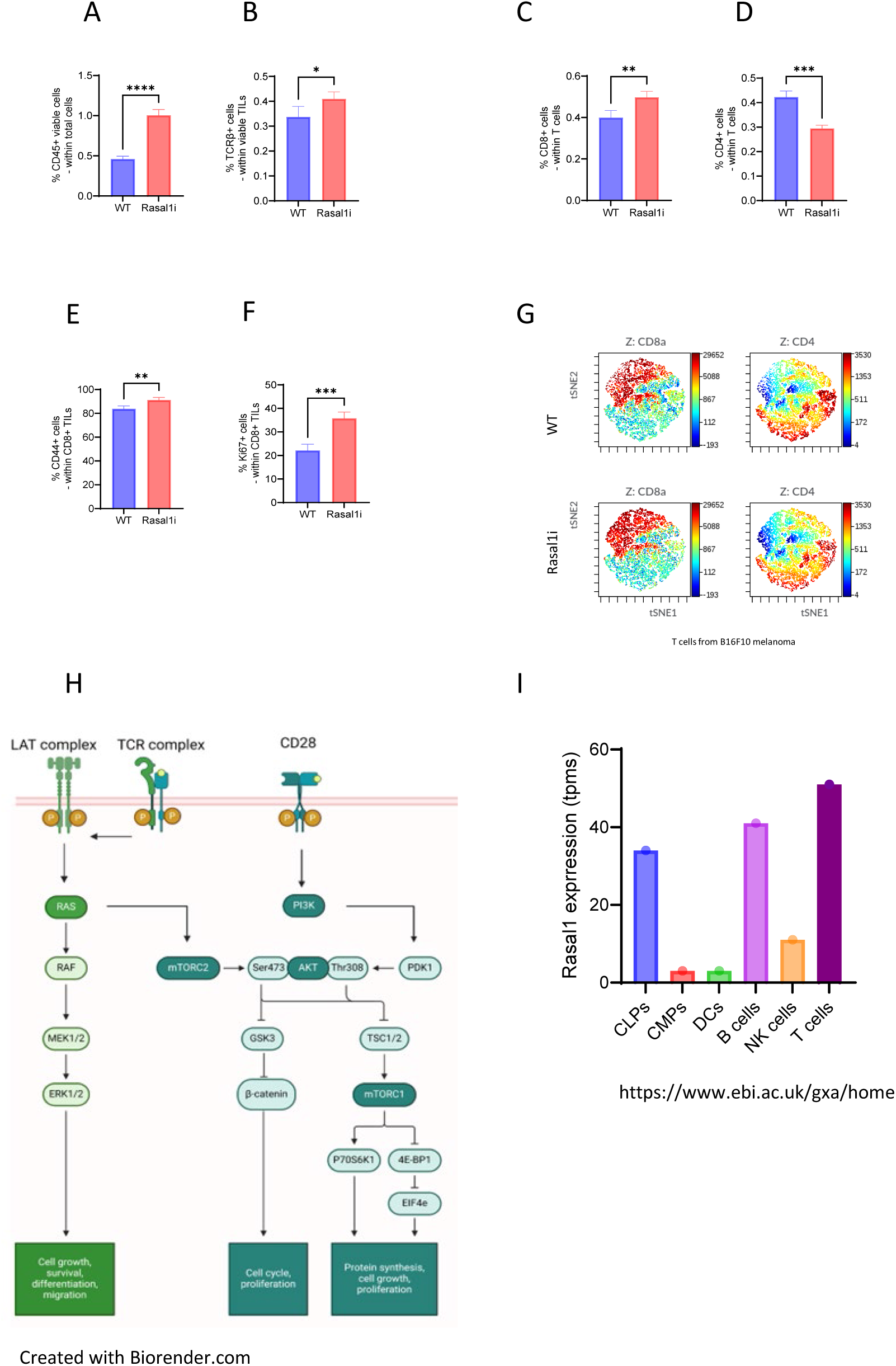
Rasal1i increases tumor-infiltrating immune cell presence in B16F10 melanoma. Percentage of viable CD45^+^ TILs detected in the tumors (**A**). Percentage of TCRβ^+^ T cells within CD45^+^ TIL population (**B**). Percentage of CD8^+^ T cells within TCRβ^+^ T population (**C**). Percentage of CD4^+^ T cells within TCRβ^+^ T cell population (**D**). Percent CD44^+^ cells within CD8^+^ TILs in melanoma (**E**). Percentage of Ki67^+^ cells within CD8^+^ TILs in melanoma (**F**). Analysis conducted on Cytobank (aka viSNE) showing the T cell compartment in the context of B16F10 melanoma (**G**). Cartoon showing the essential sequential activations occurring in T cells (**H**). Expression of Rasal1 in murine hematopoietic lineages according to the European Bioinformatics Institute database (https://ebi.ac.uk/gxa/home) (**I**). *For statistical analyses, parametric unpaired two-tailed t-tests were used. *p < 0.05, **p < 0.01 and ***p < 0.001*.

### 3.3 Rasal1i augments intratumoral conventional dendritic cell function and T cell activation

In addition to the European Bioinformatics Institute, the Immunological Genome Project (https://www.immgen.org/) also demonstrates that Rasal1 is expressed in DCs. To provide a cellular mechanism that explains the increase in the Rasal1-impaired intratumoral T cell activation, we examined the activation of intratumoral type 1 and type 2 conventional dendritic cells (cDC1s and cDC2s). We reasoned that since DCs prime T cells in the TME, this would elaborate our knowledge of Rasal1 role in T cells with respect to the activation status. First, we found that Rasal1i B16F10 bearing mice carried more DCs in the tumors when compared to the WT mice (**Fig. 3A**). In the LLC1 tumors, Rasal1i carried more DCs than WT mice (**S. Fig. 2**). DC subtypes, namely cDC1 cDC2 and mature immunoregulatory mreg DCs (PDL1^+^ DCs)[12], were found to be all similar in proportion in WT and Rasal1i mice bearing tumors (**Fig. 3A**).

**Fig. 3.**
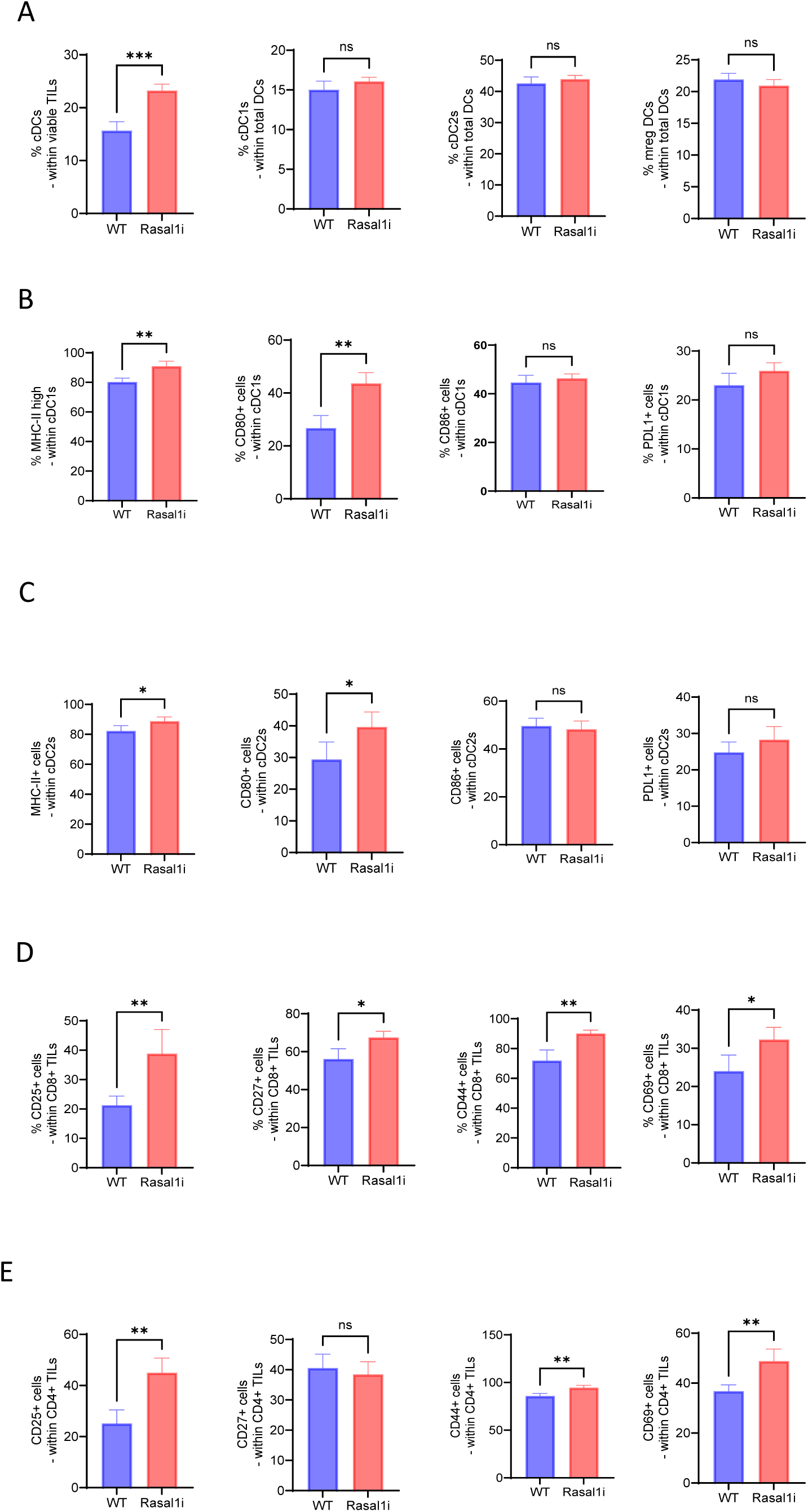
Rasal1 impairment increased DC function in B16F10 melanoma. Percentage of total cDCs, type 1 cDCs, type 2 cDCs, and mature regulatory DCs (**A**). Effect of Rasal1i on the activation of intratumoral cDCs type 1 and type 2 using the percentage of positive MHC-II, CD80, CD86, and PDL1 positivity in the context of B16F10 melanoma tumor-infiltrating Tregs (**B-C**). Rasal1i increases T cell activation in B16F10 melanoma: Percent positivity of putative activation markers (CD25, CD27, CD44 and CD69) on CD8^+^ and CD4^+^ TILs in the context B16F10 melanoma (**D-E**). *For statistical analyses, parametric two-tailed unpaired t-tests were used. *p < 0.05, **p < 0.01 and ***p < 0.001*.

cDC1s are a type of professional antigen presenting cells that specialize in cross presentation to cytotoxic CD8^+^ T cells. cDC1s (CD103^+^ DCs) express CD80 and CD86, two membrane proteins required for the interaction with the costimulatory receptor CD28 expressed on T cells. CD80 exhibits a 5-fold stronger affinity for CD28 when compared to CD86[13-15]. We found that Rasal1-impaired intratumoral cDC1s express a significant increase in MHC-II and in CD80 when compared to their WT counterparts (**Fig. 3B**). No changes were observed with respect to CD86 or for the regulatory ligand B7H1, aka PDL1 (**Fig. 3B**). Taken together, these results suggest that Rasal1 operate not only within the T cell compartment but also in the DC lineage, thereby catalyzing CD8^+^ T cell immunity to tumor antigens.

cDC2s refer to a type of professional antigen presenting cells that specialize in priming helper CD4^+^ T cells. A subset of cDC2s (CD11b^+^ DCs) induce tumor-specific CD4 anticancer Th1 responses by producing IL12[12]. Similar to cDC1s, we found that Rasal1-impaired augments the expression of MHC-II, CD80, but not that of CD86 and PDL1, on intratumoral cDC2s, when compared to their WT counterparts (**Fig. 3C**). No changes were observed with respect to CD86 or the regulatory ligand B7H1, aka PDL1 (**Fig. 3C**). Taken together, these results suggest that Rasal1 operate not only in the T cell compartment but also in the DC lineage, thereby catalyzing T cell immunity to tumor antigens.

Considering the expanding intratumoral CD8^+^ T cell compartment, and following the increased DC input to T cells, we examined the status of intratumoral T cell activation, by examining the expression of CD25, CD27, CD44, and CD69, surface receptors known to play central roles in T cell activation. Our results show a significant increase in CD25, CD27, CD44 and CD69 in response to Rasal1 impairment in tumor infiltrating T cells, except for CD27 on CD4^+^ TILs (**Fig. 3D-E**). Taken together, these results suggest that Rasal1i increases intratumoral T cell activation, with one mechanism based on interactions with DCs.

### 3.4 Rasal1i augments intratumoral T cell stemness

In addition to their central role in T cell activation, CD25, CD27, CD44 and CD69 play critical roles in shaping the formation and maintenance of memory T cells. CD25 (IL2Rα) is transiently upregulated during priming, allowing activated T cells to sense IL2 and undergo clonal expansion while also influencing the balance between terminal effector differentiation and memory precursor formation. CD27, a TNF receptor family costimulatory molecule, provides survival and anti-apoptotic signals that support the persistence of memory precursor cells and promote long-term recall capacity. CD44, an adhesion molecule and hyaluronan receptor, is stably upregulated on antigen-experienced T cells and serves as a hallmark of memory, facilitating tissue trafficking, retention, and rapid responsiveness upon antigen re-encounter. CD69, an early activation marker, contributes to memory formation by modulating tissue residency, thereby supporting the establishment of tissue-resident memory T cells. Thus, these markers are involved in memory T cell formation, survival signaling, migratory behavior, and tissue localization to orchestrate durable T cell function[16].

In order to examine the effect of the interaction between DCs and T cells, we examined the resulting TCR activation in B16F10 melanoma intratumoral T cells by quantifying the levels of phosphorylated Lck and ZAP70 at the key activation phosphorylation site Tyr 394 for Lck (pLck) and Tyr319 for ZAP70 (pZAP70), respectively. Flow cytometric analysis showed that Rasal1i CD8^+^ TILs exhibited significantly higher levels of pLck and pZAP70 in comparison to their WT counterparts (**Fig. 4A**). Since Rasal1 belongs to the GAP family, whereby it exerts its inhibitory effects on the small GTPase p21 Ras by favoring a GDP-bound form, we also reasoned to examine the effects of Rasal1i on the MAPK pathway. It has been shown that Rasal1 activity downregulates the phosphorylation of Raf, MEK and ERK in αCD3-stimulated mouse primary T cells[7]. We thus examined the effect of Rasal1i on the activation of the Ras pathway in B16F10 melanoma-derived CD8^+^ TILs. Our results showed a significant increase in the activating phosphorylation (Thr 202/Tyr 204) in the mutant CD8^+^ TILs when compared to their WT counterparts, thereby demonstrating an increased activation of the Ras pathway in CD8^+^ T cells (**Fig. 4A**). Because Ras converges onto Wnt signaling through mTORC2, Akt and GSK3, we thought to examine the activation of the Wnt pathway. For these reasons, we examined the effects of Rasal1 impairment onto pmTOR, a putative Ras effector. First, we found that Akt phosphorylation at Thr 308 was increased in Rasal1-impaired CD8^+^ melanoma TILs (**Fig. 4A**). This suggests that PI3K-Akt axis is affected in response to Rasal1 impairment. However, we found that pmTOR (Ser 2448) has been upregulated in response to Rasal1 impairment in B16F10 CD8^+^ TILs; as well as pAkt (Ser 473) thereby suggesting pmTOR complex 2 has been activated (**Fig. 4A**).

**Fig. 4.**
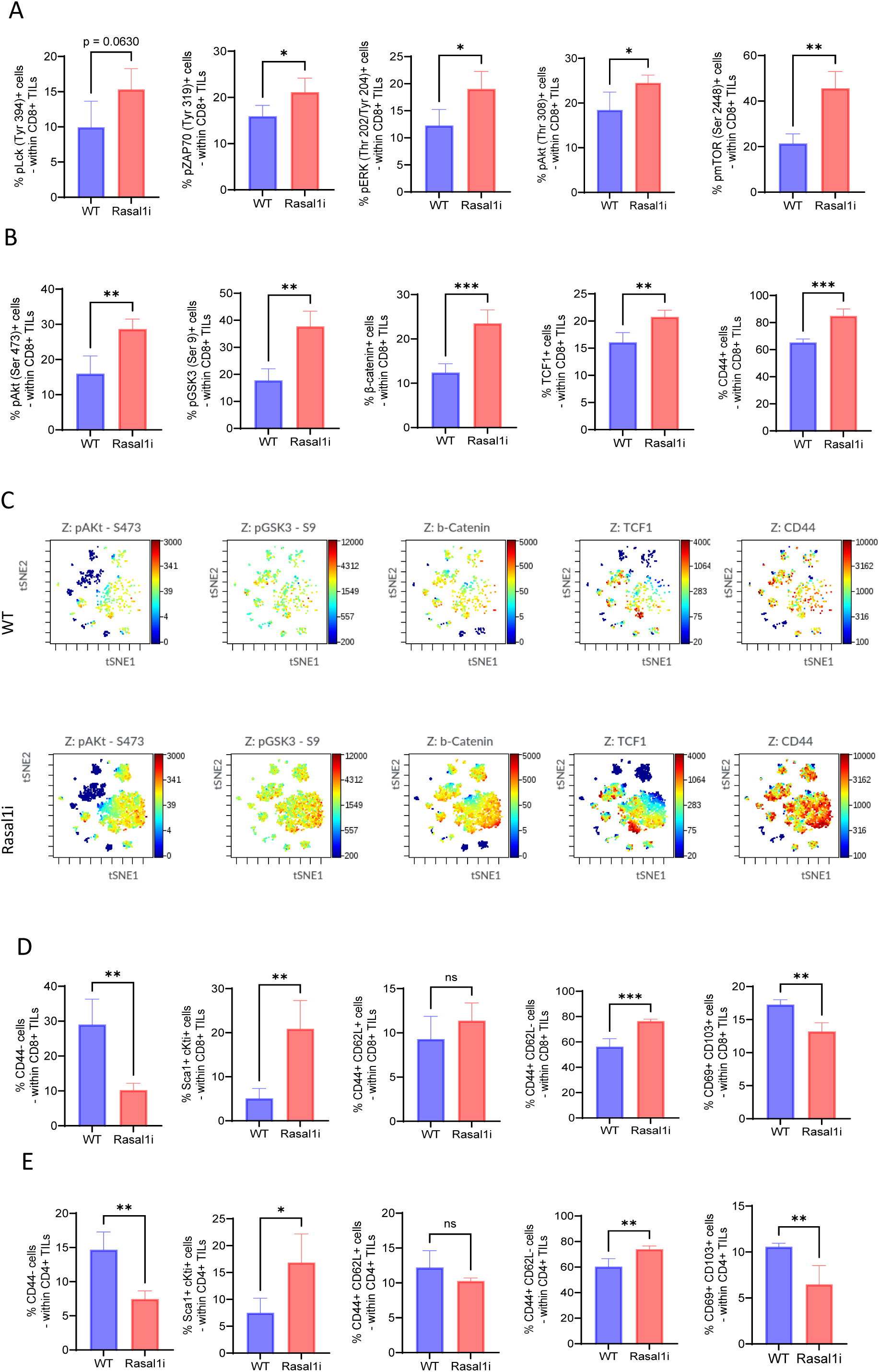
Rasal1i augments CD8^+^ TIL stemness in B16F10 melanoma. Percentage of positive pLck (Y394), pZAP70 (Y319), pERK (T202/Y204), pAkt (T308) and pmTOR (S2448) within CD8^+^ TILs in melanoma (**A**). Percentage of positive pAkt (S473), pGSK3β (S9), β-catenin, TCF1 and CD44 within CD8^+^ TILs in melanoma (**B**). A proportional viSNE analysis conducted on Cytobank showing the colocalization of Wnt signaling active members within the CD8^+^ TIL compartment in the context of B16F10 melanoma (**C**). Percentage values of putative memory subsets in CD8^+^ TILs in the context B16F10 melanoma (**D**). Percentage values of putative memory subsets in CD4^+^ TILs in the context B16F10 melanoma (**E**). *For statistical analyses, parametric unpaired two-tailed t-tests were used. *p < 0.05, **p < 0.01 and ***p < 0.001*.

In light of the observed increased Ras and thereby T cell activation, and given crosstalk of the Ras pathway with stem cell maintenance pathways, we reasoned to examine the Rasal1i ability to regulate T cell progenitors. We also examined the effects of Rasal1i on the Wnt pathway in CD8^+^ TILs. When compared to the WT, Rasal1-impaired CD8^+^ TILs demonstrated a significant increase in pAkt (Ser 473), pGSK3β (Ser 9), β-catenin, TCF1, and CD44 (**Fig. 4B**). A viSNE analysis demonstrated that pGSK3β^+^ cells colocalized with pAkt^+^ cells, β-catenin^+^ cells, TCF1^+^ cells, and CD44^+^ cells thereby demonstrating the unleashing of the Wnt pathway in the Rasal1-impaired CD8^+^ TILs (**Fig. 4C**). These data suggest that Rasal1i upregulates the Wnt pathway in CD8^+^ TILs, induces their expansion, and expands the stem-like cell reservoir in the tumor.

Memory T cells are crucial cells in the maintenance of an immune response against cancer, whereby one hypothesis postulates that TCR signal strength stimulates the formation of memory T cells[17]. Upon TCR ligation, the Ras pathway is rapidly activated through LAT; and mice lacking the N-Ras isoform were found capable of mounting a CD8^+^ T cell effector phenotype against viral pathogens but failed to generate protective memory cell. This result highlights a key role of Ras in the formation of memory CD8^+^ T cells[18]. We thus examined the effects of Rasal1i on the frequency of memory CD4^+^ TILs in the context of B16F10 melanoma. Rasal1i resulted in several changes within the infiltrating memory T cell compartment (**Fig. 4D-E**). Naïve like CD44^-^ T cells were decreased for both CD4^+^ and CD8^+^ TILs; stem-like memory T cells Sca1^+^ cKit^+^ were increased for both CD4 and CD8^+^ TILs; central memory CD44^+^ CD62L^+^ were similar for both CD4^+^ and CD8^+^ TILs; effector memory CD44^+^ CD62L^-^ were increased for both CD4^+^ and CD8^+^ TILs; and resident memory CD69^+^ CD103^+^ were decreased for both CD4^+^ and CD8^+^ TILs in response to Rasal1 impairment in B16F10 melanoma. However, it is important to note that despite these changes in the frequency of memory T cells, the total numbers of all subsets of T memory cells in the tumor were significantly increased in the Rasal1i mice due to the increased infiltration, suggesting an overall infiltration of TILs regardless of the subset. These results were corroborated in MC38 CD8+ TILs and LLC1 CD4+ TILs (**S. Fig. 2C-D**). Taken together, these results suggest that Rasal1i increases T cell stemness in cancer.

### 3.5 Rasal1i promotes anticancer T cell function

T-bet drives effector differentiation, cytotoxicity, and IFN-γ production in CD8^+^ TILs against cancer, whereas CXCR5 marks a distinct subset of CD8^+^ T cells with stem-like, progenitor-exhausted properties. These cells retain self-renewal capacity, localize to lymphoid niches within tumors, and serve as the main pool that expands in response to immune checkpoint blockade. For this reason, we then analyzed how CD8^+^ TILs contributed to tumor control, by factoring in tumor reactivity as indicated by Tbet, PD1 expression, CXCR5, and granzyme B. Rasal1 impairment caused a significant increase of Tbet, PD1, CXCR5, and granzyme B in CD8^+^ TILs (**Fig. 5A**). Helper CD4^+^ TIL analysis showed a significant increase in the expression of Tbet in response to Rasal1 impairment when compared to WT mice (**Fig. 5B**). Furthermore, helper CD4^+^ TILs showed an increased expression of CXCR5 and PD1, but similar level of granzyme B in Rasal1i mice as compared to WT (**Fig. 5B**). viSNE analysis demonstrated how Rasal1i resulted in colocalization of increased Tet, CXCR5, and PD1 in CD8^+^ TILs (**Fig. 5C**). Furthermore, in CD4^+^ T cell subsets, we found an inverse pattern between the expression of Rasal1 and Tbet, and a weaker pattern between Rasal1 and PD1 (https://www.ebi.ac.uk/gxa/home) (**Fig. 5D**). Altogether, these results suggest that Rasal1 impairment augments anticancer T cell cytolytic response.

**Fig. 5.**
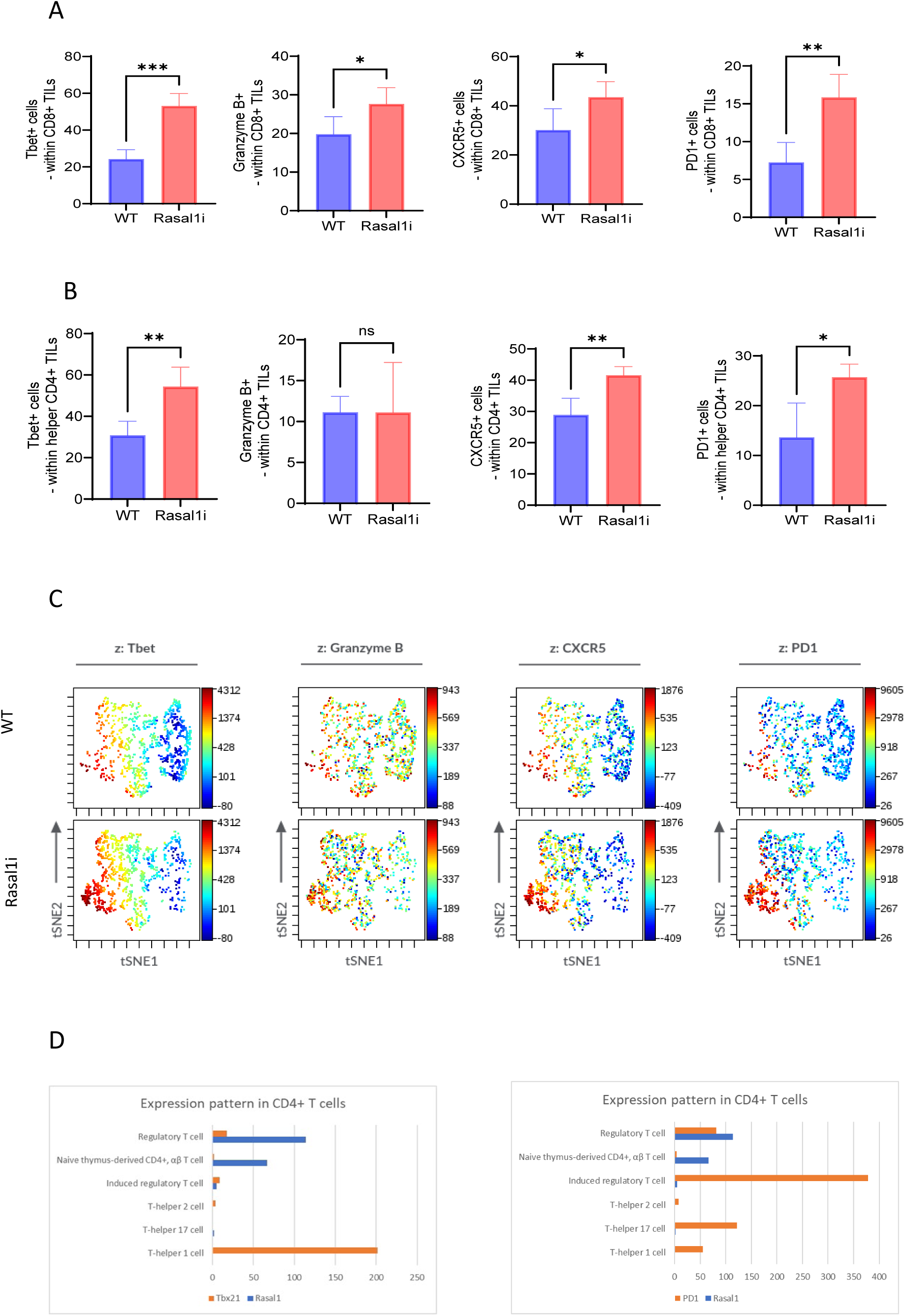
Rasal1i increases tumor-infiltrating T cell cytolytic activity against B16F10 melanoma. Percentage of Tbet^+^, GZB^+^, CXCR5^+^ and PD1^+^ cells within CD8^+^ TILs in B16F10 melanoma (**A**). Percentage of Tbet^+^, CXCR5^+^ and GZB^+^ cells within helper CD4^+^ TILs (Foxp3^-^ cells) in B16F10 melanoma (**B**). A viSNE analysis conducted on Cytobank showing the colocalization of Tbet, GZB, CXCR5 and PD1 within the CD8^+^ TIL compartment in the context of B16F10 melanoma (**C**). Comparison of relative expression values between Rasal1 expression versus T-bet (left) and PD1 (right) in murine CD4^+^ T cell lineages (data collected from the the European Bioinformatics Institute database; **https://ebi.ac.uk/gxa/home**) (**D**). *For statistical analyses, parametric unpaired two-tailed t-tests were used. *p < 0.05, **p < 0.01 and ***p < 0.001*.

### 3.6 Rasal1i overrides PDL1 negative signaling and potentiate αPD1 in B16F10-PDL1 melanoma

Given that Rasal1 acts as intracellular checkpoint that negatively modulates CD8^+^ TIL responses, we next examined the effects of Rasal1 impairment on B16F10 tumors. The idea behind this approach is that the “intracellular” or “downstream” characteristic of Rasal1 impairment overrides the negative signaling incoming from surface checkpoints such as PD1. For this reason, we employed the PDL1-expressing B16F10 cancer cell in the following experiment and examined the effects of Rasal1 impairment on survival in PDL1-overexpressing B16F10 tumors in females. Rasal1 impairment prolonged survival in B16F10-PDL1 melanoma in females (**Fig. 6A**). Specifically, by day 16 post-implantation in females, WT mice started to display disease symptoms, including pale palms and slower movements, whereas mutant mice appeared healthier, and capable of performing their physiological activities until day 22 post implantation, when the last mouse was sacrificed. Of note, we employed intradermal skin implantation, a model in which skin bleeding from the tumor would cause a rapid death to the mouse. We believe that a 4-day advantage is compelling evidence for survival prolongation given the intradermal nature of the melanoma implantation we employed (**Fig. 6A**). Furthermore, and for this purpose, we performed αPD1 in WT and Rasal1-impaired mice. WT and Rasal1i mice were implanted with B16F10-PDL1 melanoma cells intradermally. The injections of αPD1 (100 µg/mouse) were started at day 4 post implantation, every other day throughout the experiment. Rasal1 impaired mice showed a synergistic effect in the B16F10 melanoma that overexpressed PDL1 (**Fig. 6B**). Statistical analyses showed that αPD1 and Rasal1 mutants AUC were significantly lower than WT. Combo AUC was significantly lower than any other treatment, whereas αPD1 was similar Rasal1 mutant throughout the experiment (**Fig. 6B**).

**Fig. 6.**
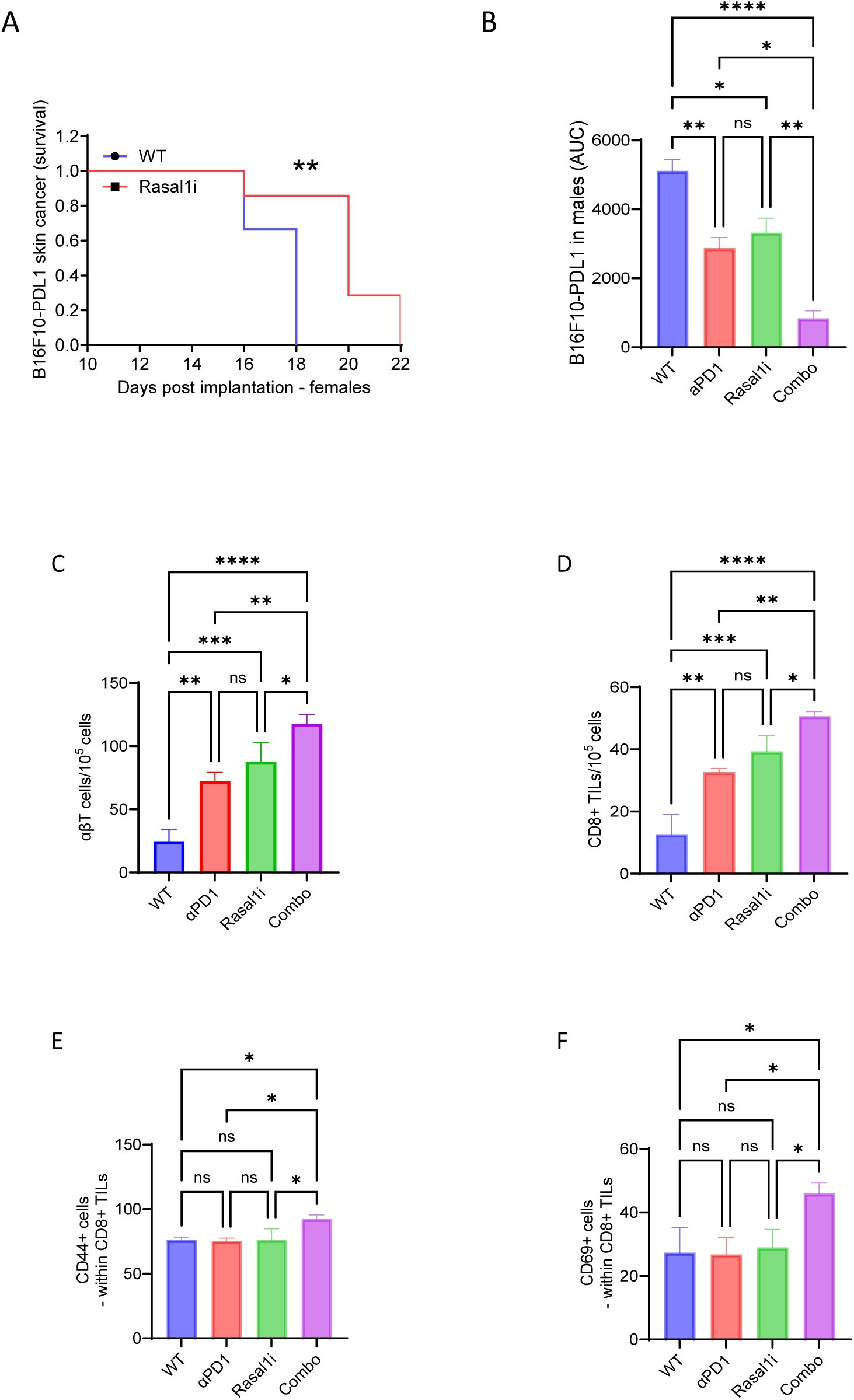
Rasal1i potentiates *α*PD1 in resistant in B16F10 melanoma. Survival in females and AUC in males derived from intradermal implantation of B16F10-PDL1 cells (**A-B**). Counts of infiltrating T cells and CD8^+^ T cells within B16F10-PDL1 melanoma (**C-D**). Percent CD44^+^ and CD69^+^ cells within gated effector CD8^+^ TILs in B16F10-PDL1 melanoma (**E-F**). *For statistical analyses, parametric one-way ANOVAs were used, with multiple comparisons. *p < 0.05, **p < 0.01 and ***p < 0.001*.

When we phenotyped the B16F10-PDL1 TILs, we found that αPD1, Rasal1 mutants, and Combo carried significantly more total T cells as well as CD8^+^ T cells in their tumors as compared to WT (**Fig. 6C-D**). Interestingly, we found that the percentages of CD44^+^ cells, as well as of CD69^+^ cells within intratumoral T cells, were significantly higher in the Combo group as compared to any other group (**Fig. 6E-F**). Taken together, our results show that Rasal1 impairment overrides the PDL1 negative signaling imposed by the TME, partially by means of increasing T cell infiltration into the tumors; and that Rasal1 impairment potentiates αPD1 ICB in melanoma.

### 3.7 RASAL1 is upregulated in tumors but shows limited prognostic association in bulk TCGA cohorts

To assess the clinical relevance of RASAL1 in modulating anti-tumor immunity, we examined its expression and prognostic impact in TCGA cohorts for melanoma (SKCM), colorectal cancer (COAD/READ), and non-small cell lung cancer (LUAD/LUSC). RASAL1 mRNA levels were significantly higher in tumor versus normal tissue in colorectal cancer (Wilcoxon *p*=0.0004; *n*=647 tumors, *n*=51 normals) and lung cancer (*p*<2.22e-16; *n*=1051 tumors, *n*=110 normals) (**Fig. 7A**). In melanoma, expression was numerically higher in tumors but not significant (*p*=0.97; *n*=103 tumors, *n*=1 normal), likely due to the extremely limited number of matched normal samples in TCGA-SKCM (**Fig. 7A**).

**Fig. 7.**
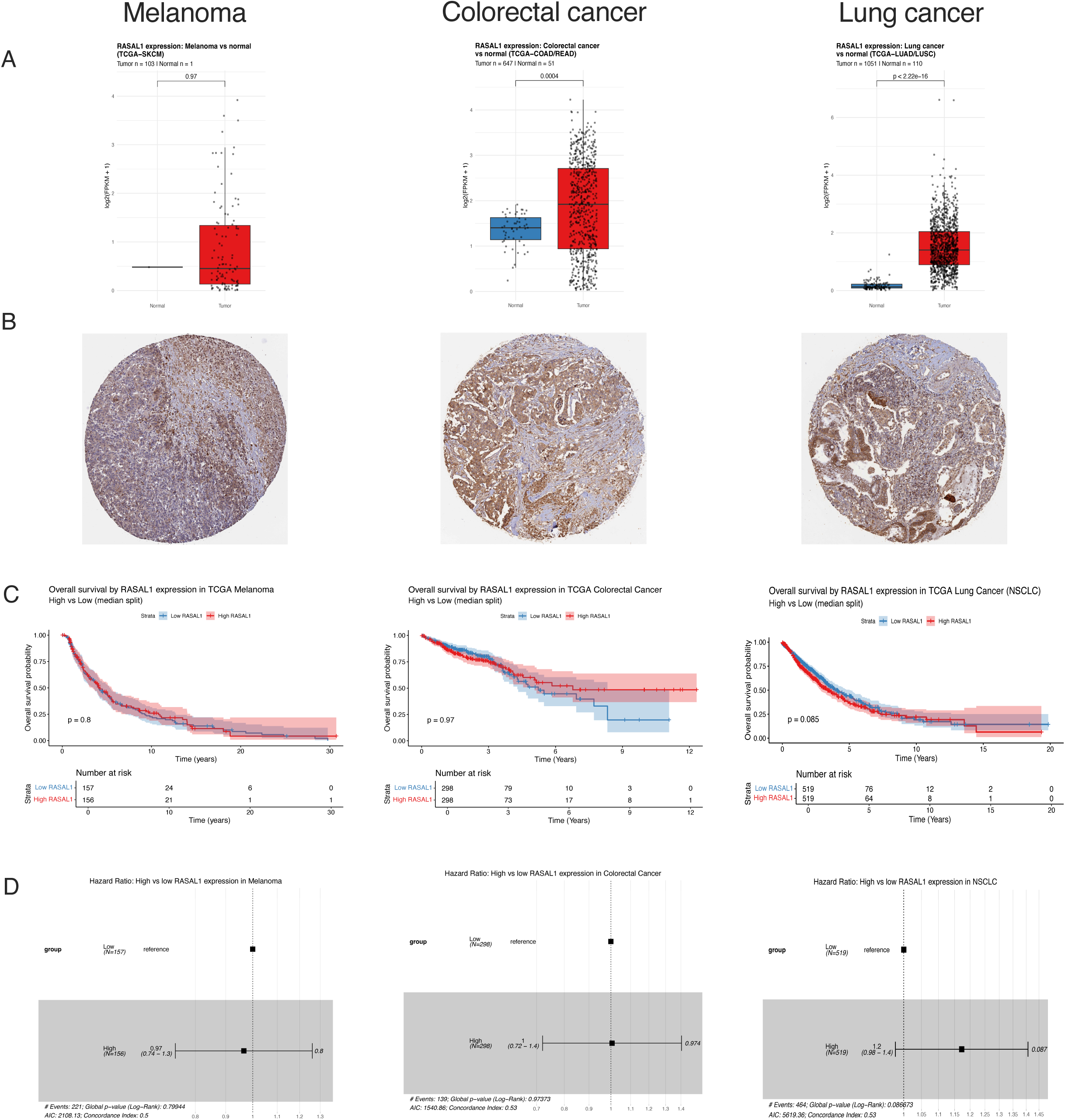
RASAL1 expression and prognostic associations in TCGA melanoma, colorectal cancer, and non-small cell lung cancer cohorts. (**A**) Boxplots of log₂(FPKM+1) RASAL1 expression in tumor vs normal tissue. Top: melanoma (TCGA-SKCM; *n*=103 tumors, *n*=1 normal; Wilcoxon *p*=0.97; note: severely limited normal samples preclude robust comparison). Middle: colorectal adenocarcinoma (TCGA-COAD/READ; *n*=647 tumors, *n*=51 normals; *p*=0.0004). Bottom: non-small cell lung cancer (TCGA-LUAD/LUSC; *n*=1051 tumors, *n*=110 normals; *p*<2.22e-16). Red = tumor, blue = normal. Horizontal lines: median; whiskers: 1.5× interquartile range; points: individual samples. (**B**) Representative immunohistochemistry images of RASAL1 protein expression in punch biopsies from the Human Protein Atlas (HPA). Top: malignant melanoma (metastatic site) showing low staining, moderate to weak intensity, <25% quantity, cytoplasmic/membranous/nuclear localization. Middle: colon adenocarcinoma showing medium staining, moderate intensity, >75% quantity, cytoplasmic/membranous. Bottom: lung adenocarcinoma showing medium staining, moderate intensity, >75% quantity, cytoplasmic/membranous. Images selected as representative of the most common/average patterns reported in HPA for each cancer type. (**C**) Kaplan–Meier survival curves stratified by median RASAL1 expression (log₂(FPKM+1)). Top: melanoma (TCGA-SKCM; High *n*=156 red vs Low *n*=157 blue; log-rank *p*=0.8). Middle: colorectal (TCGA-COAD/READ; High *n*=298 red vs Low *n*=298 blue; *p*=0.97). Bottom: NSCLC (TCGA-LUAD/LUSC; High *n*=519 red vs Low *n*=519 blue; *p*=0.085). Shaded areas: 95% CI. Risk tables below each plot show patients at risk over time. (**D**) Forest plots from univariate Cox regression. HR compares high vs low RASAL1 (low = reference). Top: Melanoma (*n*=313; HR=0.97 [0.74–1.3]). Middle: Colorectal (*n*=596; HR=1 [0.72–1.4]). Bottom: NSCLC (*n*=1096; HR=1.2 [0.98–1.4]). Error bars: 95% CI.

Representative immunohistochemistry from the Human Protein Atlas (HPA) revealed corresponding protein patterns in tumor punch biopsies. Melanoma (metastatic site) displayed low staining with moderate to weak intensity, less than 25% positive tumor cells, primarily cytoplasmic/membranous with occasional nuclear positivity in some cases (**Fig. 7B**). In contrast, colorectal adenocarcinoma showed medium staining of moderate intensity with greater than 75% positive cells, confined to cytoplasmic/membranous compartments (**Fig. 7B**). Lung adenocarcinoma exhibited a similar profile with medium staining, moderate intensity, and greater than 75% positivity in cytoplasmic/membranous regions (**Fig. 7B**). In bulk tumor survival analyses (median split), Kaplan–Meier curves showed no significant OS differences between high- and low-RASAL1 groups in melanoma (log-rank *p*=0.8) or colorectal cancer (*p*=0.97), with only a borderline trend toward poorer survival in high-expression lung cancer cases (*p*=0.085) (**Fig. 7C**). Univariate Cox regression confirmed no significant risk alterations (HR ≈1, 95% CI crossing 1; melanoma HR=0.97 [0.74–1.3], colorectal HR=1 [0.72–1.4], lung HR=1.2 [0.98–1.4]) (**Fig. 7D**).

Overall, these findings indicate RASAL1 upregulation in bulk colorectal and lung tumors (likely tumor-intrinsic), with weak or absent prognostic signals, consistent with dominant tumor-cell expression overshadowing potential contributions from immune infiltrates.

## 4. DISCUSSION

A visible proportion of cancer patients do not respond to current immunotherapies, thereby necessitating an urgent need for the identification of novel inhibitory checkpoints. Intracellular inhibitory checkpoints are advantageous over cell surface ones since tumoral cells cannot access the intracellular “breaks” in NK or T cells in similar manner to PDL1 and/or PD1 interaction. Thus, the removal of intracellular immune checkpoints from effector immune cells bypasses inhibitory signaling, promotes activation of effector immune cells, thereby catalyzing the elimination of target cells^9^.

GTPase-activating protein (GAP) Rasal1, acts upon the central GTPase Ras to inhibit its activity, was characterized as an intracellular immune checkpoint that negatively regulates T cell function[7]. Rasal1 was also expressed at low levels in naïve T cells and was upregulated following stimulation of both CD4^+^ and CD8^+^ T cells, thereby revealing the behavior of an immune checkpoint. The GTPase activity of p21 Ras acts as a molecular switch to activate RAF upon the hydrolysis of GTP to GDP, thereby activating the MAPK pathway. Activated RAF phosphorylates MEKs 1&2, which in turn phosphorylate ERKs 1&2. Activated ERKs translocate to the nucleus whereby they modulate the activity of transcriptions factors involved in cell proliferation, differentiation, and effector functions, thereby translating the activation of the MAPK signaling pathway. Rasal1 inhibited the MAPK signaling by blocking p21 Ras function. Rasal1 knockdown significantly reduced B16F10 melanoma and EL4 lymphoma tumors, highlighting the therapeutic potential of Rasal1 for anticancer immunotherapy[7].

The fact that Rasal1 is an intracellular molecule is very advantageous. This characteristic renders effector immune cells continuously effective in eliminating tumoral cells despite PDL1-PD1 interactions. Because Ras is downstream of the TCR and PD1, the inhibition of Rasal1 and thereby the unleashing of Ras, overrides the inhibitory signaling incoming from T cell surface. The inaccessibility of Ras signaling axis to the tumoral cell endows the cytotoxic CD8^+^ T cells with the capability to escape inhibition by resistance mechanisms applied by the tumoral cells, or possibly by Treg cells. Rasal1i induced a significant B16F10 and B16F10-PDL1 melanoma control. Rasal1 impairment also mitigated the growth of MC38 colorectal cancer, and that of the “cold” tumor LLC1 lung cancer[9]. These results have several implications in which the intracellular nature of Rasal1 did indeed override the inhibitory signaling due to PDL1 in melanoma and did indeed enabled immunity against “cold” tumors such as LLC1 lung cancer.

The tumor control introduced by Rasal1i appears to have to resulted in part from the expansion of the CD8^+^ TIL compartment, and its cytolytic activity. The alteration observed in the T cell compartment in response to Rasal1i suggests that Rasal1 plays a central role in the expansion of effector immune cells inside the tumor. We observed that Ras pathway has been unleashed in the CD8^+^ TILs. The observation that a visible proportion of cells demonstrated positivity for Ki67 in the WT CD8^+^ TILs, combined with the observation that Ki67 positivity was significantly increased in Rasal1 mutant CD8^+^ TILs, suggest that Rasal1 directly regulates cell cycle intratumorally in response to neoantigens. This was corroborated by the significant increases we observed in the activation markers CD25, CD27, CD69 and CD44.

Rasal1 impairment expanded the reservoir of CD8^+^ TIL in a very powerful manner. For this reason, we thought to examine the CD8 progenitor compartment. Rasal1 impairment promoted the generation of stem-like CD8^+^ T cells. This discovery is of significant implication to cancer immunotherapy. Tscm refer to a subset of memory T cells that possess stem cell-like properties, including self-renewal and the ability to differentiate into other memory and effector T cell subsets. Tscm are characterized by the expression of certain surface markers Sca1 and cKit (aka CD117), which distinguish them from other memory T cell subsets. These cells are thought to play a crucial role in long-term immune responses, as they can persist for extended periods and give rise to other memory T cell subsets upon encountering antigen[19]. Tscm have been identified in various contexts, including viral infections[20], autoimmune diseases, and cancer, where they may contribute to combatting disease pathogenesis. The significant increase in Tscm within tumors could have several implications. First, Tscm are believed to possess superior proliferative and self-renewal capacities compared to other T cell subsets. Their increase within the tumor suggests a potential for long-lasting immune responses against the tumor. Second, Tscm have been proposed as a promising target for immunotherapy due to their stem cell-like properties. Their increased abundance within the tumor could indicate a favorable environment for immunotherapy, such as adoptive T cell transfer or checkpoint blockade, to be effective. Third, the presence of Tscm within the tumor could serve as a favorable prognostic marker for patient outcomes. Higher levels of Tscm might correlate with improved survival or response to therapy in certain cancers. Fourth, this discovery highlights the importance of further research into the role of Tscm in tumor immunity. Understanding the mechanisms that regulate Tscm function within tumors could lead to the development of novel therapeutic strategies targeting these cells[21].

The canonical Wnt signaling pathway is comprised of a family of secreted signaling modulators that bind to transmembrane G-protein coupled receptors termed Frizzled. Frizzled receptors interact with the LRP 5&6 coreceptors, which recruit the cytoplasmic member Dishevelled. In an off-state, Dishevelled roams the cytoplasm, and allows the destruction complex, which is composed of Axin, APC, CK1 and GSK3, to phosphorylate β-catenin and thereby target it for proteasomal degradation. In an on state, Dishevelled is recruited to the cytoplasmic membrane whereby it anchors the destruction complex to the plasma membrane, thereby liberating β-catenin. Then, β-catenin translocates to the nucleus whereby it associates with other transcription factors such as TCF1 to alter gene expression. GSK3 inhibition has been used as a therapeutic strategy to activate the Wnt signaling for regenerative medicine and stem cell expansion. Further, the Wnt pathway has been demonstrated to play key roles in the maintenance and regulation of CD8^+^ T cell progenitors and CD8^+^ memory T cells[22]. Furthermore, it has also been well established that activated ERK phosphorylates GSK3 and renders it inactivated in T cells[23-27], suggesting that activated ERK contributes to the activation of the Wnt pathway. Wnt signaling activation causes an increase in memory TILs presence[28]. In our work, Rasal1i caused a significant increase in Wnt pathway activation and promoted TCF1 in CD8^+^ T cells. The significant and substantial increase in Wnt signaling in intratumoral CD8^+^ T cells could have several implications. First, Wnt signaling is known to promote T cell survival and proliferation. The increased Wnt signaling in intratumoral CD8^+^ T cells may indicate a mechanism by which these cells are sustained and expanded within the tumor microenvironment. Second, activation of the Wnt signaling pathway in CD8^+^ T cells has been associated with enhanced effector function and antitumor activity. The increased Wnt signaling in intratumoral CD8^+^ T cells could therefore contribute to a more effective immune response against the tumor. Third, Wnt signaling has been implicated in the regulation of T cell exhaustion, a state of functional impairment that occurs in chronically stimulated T cells within the tumor microenvironment. Increased Wnt signaling in intratumoral CD8^+^ T cells may help protect these cells from exhaustion, maintaining their ability to target and eliminate tumor cells. Fourth, Wnt signaling pathway is a druggable target, and its modulation has been explored in cancer therapy. The increased Wnt signaling in intratumoral CD8^+^ T cells suggests that targeting this pathway could be a strategy to enhance the efficacy of immunotherapy in cancer. Fifth, the level of Wnt signaling in intratumoral CD8^+^ T cells could serve as a biomarker for predicting response to immunotherapy. Patients with higher Wnt signaling in their intratumoral CD8^+^ T cells may be more likely to respond to immune checkpoint blockade or other immunotherapies. Overall, these implications highlight the potential significance of increased Wnt signaling in intratumoral CD8^+^ T cells and suggest avenues for further research and therapeutic development in cancer immunotherapy[29,28,30].

The specific mechanism responsible for driving this proliferation remained elusive until Siddiqui et.al. (2019) described the role of TCF1 in mediating such effects[31]. Siddiqui identified a subset of CD8^+^ TILs that express both PD1 and TCF1 that are central to the response to immunotherapy. The absence of these TCF1^+^ PD1^+^ CD8^+^ TILs limit the response to immunotherapy. Importantly, TCF1 was found essential for their stem-like T cells. Siddiqui and colleagues postulated that “similar cells were found in human melanoma patients, indicating that the success of immune checkpoint blockade doesn’t stem from reversing T cell exhaustion but rather from harnessing the proliferative capacity of these stem-like T cells”. As demonstrated by Siddiqui et al. (2019), these cells are responsible for tumor control in immune checkpoint blockade. The progenitors constitute a source that produce more of both TCF1^+^ progenitors and TCF1^-^ CD8^+^ TILs. Our results align with the findings of the study conducted by Siddiqui and colleagues[31]. Indeed, Rasal1 impairment resulted in a significant increase in intratumoral CD44^+^, CD69^+^ CD8^+^ TILs. These results imply that Rasal1 impairment results in the expansion of the whole CD8^+^ TIL compartment rather than a specific compartment, thereby promoting tumor control.

The increase in T-bet, Granzyme B, CXCR5, and PD-1 expression in CD8^+^ tumor-infiltrating T cells following Rasal1 inhibition reveals a state of enhanced effector differentiation coupled to immune checkpoint engagement. Furthermore, the concomitant increase in T-bet, GZB, CXCR5 and PD1 on CD8^+^ T cells suggests a potential for these cells to undergo chemotaxis to regions rich with CXCL13, a chemokine that attracts CXCR5^+^ cells. Our literature search indicates that melanoma cells are rich in CXCL13[32-34]. Elevated T-bet and Granzyme B indicate robust type 1 polarization and cytotoxic potential, consistent with heightened antitumor activity. At the same time, the upregulation of PD1 suggests that these activated CD8^+^ T cells are subjected to adaptive inhibitory feedback within the tumor microenvironment, a hallmark of antigen-experienced effector cells encountering chronic stimulation. Together, this phenotype suggests that Rasal1 inhibition effectively drives CD8^+^ T cells into a highly activated, tumor-reactive state, while simultaneously creating a dependency on PD1-mediated inhibition that limits their full effector function[35].

The strong inverse correlation between Rasal1 expression and T-bet levels in CD4^+^ T cells indicates that Rasal1 acts as an intracellular restraint on Th1 differentiation within the tumor microenvironment. Given the central role of T-bet in promoting type 1 helper functions and supporting cytotoxic antitumor immunity, elevated Rasal1 expression likely limits CD4^+^ T cell effector polarization and helper capacity. In contrast, the weak association between Rasal1 and PD1 expression suggests that Rasal1-mediated regulation occurs largely independently of canonical immune checkpoint pathways. Rather than directly enforcing exhaustion, Rasal1 appears to modulate upstream activation and lineage commitment, shaping CD4^+^ T cell functional identity prior to overt checkpoint engagement. These findings support a model in which Rasal1 inhibition enhances Th1 polarization, while PD1 blockade sustains downstream effector function, providing a complementary rationale for combination therapy[36,37,35].

This provided a strong mechanistic rationale for combining Rasal1 inhibition with αPD1 therapy. Consistent with this model, we observed a clear synergy between Rasal1 inhibition and αPD1 treatment, supporting the concept that Rasal1 functions as an intracellular immune checkpoint whose inhibition sensitizes T cells to immune checkpoint blockade and amplifies therapeutic efficacy.

Rasal1i in T cells markedly enhanced stemness, effector function, and tumor control in our mouse models, yet bulk TCGA analyses across melanoma, colorectal cancer, and NSCLC show no robust prognostic association with RASAL1 expression. This is complemented by differential expression showing significant RASAL1 mRNA upregulation in colorectal and lung tumors versus normal tissue, and numerically higher (non-significant) levels in melanoma (limited by *n*=1 normal). Protein-level IHC from the HPA further supports this, with higher positivity in colorectal and lung adenocarcinomas compared to low expression in melanoma. These patterns underscore RASAL1’s divergent, context-dependent roles: inhibitory in T cells via the MAPK axis, where its loss boosts anti-tumor immunity, versus its loss in cancer cells may tumorigenic. Upregulation in bulk colorectal and lung tumors likely reflects tumor-intrinsic adaptation to try to shut down RAS signaling in response to exaggerated proliferation. Bulk RNA-seq inherently prioritizes abundant tumor signals over sparse T-cell infiltrates, explaining the weak prognostic associations despite clear expression differences. Melanoma data remain limited by scarce normals but align with the overall trend. Collectively, these results position RASAL1 as a highly context-specific regulator and support selective T cell-targeted inhibition (e.g., CRISPR editing or small-molecule approaches in adoptive therapies) to enhance anti-tumor immunity in these cancers without exacerbating tumor-intrinsic RASAL1 upregulation.

## 5. CONCLUSION

Overall, this study demonstrates a novel role of in negatively regulating T cell stemness and anticancer immunity. Rasal1 inhibition significantly mitigated anticancer responses against several cancers, and we are looking to develop small molecule inhibitors to target Rasal1. We believe that targeting Rasal1 in the context of T cells could be a novel strategy in targeting a variety of cancers.

## 6. STUDY LIMITATIONS

Given the fact that mouse model that has been used in this study is a global Rasal1 impaired model, other cells different such as non-immune cells are also contributing the anticancer effects observed. That said, we believe that a robust anticancer effect necessitates a collaboration between a multitude of immune lineages to drive an antitumoral response. These lineages could include hematopoietic ones and mesenchymal and tissue specific.

## Abbreviations

AUC: area under the curve
ICB: immune checkpoint blockade
Ras: rat sarcoma
GAP: GTPase-activating protein
RNA-seq: RNA sequencing
DCs: dendritic cells
MAPK: mitogen-activated protein kinase
RAF: rapidly accelerated fibrosarcoma
MEK: MAPK/ERK kinase
ERK: extracellular signal-regulated kinase
GDP: guanosine diphosphate
TCR: T cell receptor
PD-1: programmed cell death protein 1
PD-L1: programmed death-ligand 1
CD8^+^: cluster of differentiation 8-positive
CD4^+^: cluster of differentiation 4-positive
TILs: tumor-infiltrating lymphocytes
Ki67: marker of proliferation Ki-67
Myc: MYC proto-oncogene
T-SCM: stem-like memory T cells
Wnt: Wingless-related integration site
CLPs: common lymphoid progenitors
LRP5/6: low-density lipoprotein receptor-related protein 5/6
APC: adenomatous polyposis coli
CK1: casein kinase 1
GSK3: glycogen synthase kinase 3
TCF1: T cell factor 1
CXCR5: C-X-C chemokine receptor type 5
CXCL13: C-X-C motif chemokine ligand 13
TCGA: The Cancer Genome Atlas
NSCLC: non-small cell lung cancer.

## Acknowledgments

None.

## Funding

This work was funded by the Canadian Institutes of Health Research.

## Author contributions

MEI: conceptualization, mouse work, first manuscript draft, figure preparation; ASC: bioinformatics, revision of the manuscript, figure preparation. Both authors have approved the final version of the manuscript.

## Supplementary Figure legends

**S. Fig. 1. Rasal1i augments T cell activation and proliferation *in vitro*.** T cell counts and CFSE proliferation histograms in CD8^+^ T cells following αCD3ε/αCD28 72h or 48h stimulation *in vitro*, respectively (**A**). CD44 and CD69 activation markers in CD8^+^ T cells following in vitro 48h stimulation with αCD3ε/αCD28 (**B**). CTLA4 and PD1 deactivation markers in CD8^+^ T cells following in vitro 48h stimulation with αCD3ε/αCD28 (**C**). Fold change in mRNA expression of selected cytokines in T cells following αCD3ε/αCD28 48h stimulation *in vitro* (**D**). Effect of IL1b, IL6 and TNFa on CD8^+^ and CD4^+^ T cells based on the immunological genome project database (immgen.org) (**E**). *For statistical analyses, parametric unpaired two-tailed t-tests were used. *p < 0.05, **p < 0.01 and ***p < 0.001*.

**S. Fig. 2. Rasal1i increases immune cell infiltration in cancer and modulates markers of stem-like memory T cells in lung and colorectal cancers.** Percent viable CD45^+^ cells and DCs in lung cancer (**A**). Percent viable CD45^+^ and CD8^+^ T cells in colorectal cancer (**B**). MFI values of memory markers CD44, CD62L, Sca1 and cKit (aka CD117) in CD4^+^ TILs in LLC1 lung cancer (**D**). MFI values of memory markers CD44, CD62L, Sca1 and CD127 in CD8^+^ TILs MC38 colorectal cancer (D). *For statistical analyses, parametric unpaired two-tailed t-tests were used. *p < 0.05, **p < 0.01 and ***p < 0.001*.

## Supplementary Table legends

**S. Table 1. List of antibodies and chemicals used for this work.**

**S. Table 2. List and sequences of primers used for this work.**

**S. Table 3. Gating strategies for immune cells in this work.**

## Notes

### Competing Interest Statement

The authors have declared no competing interest.

