## Supplementary figures and images for "Rasal1 impairment unleashes anticancer immunity - a focus on T cells"

### Supplemental Figure 1

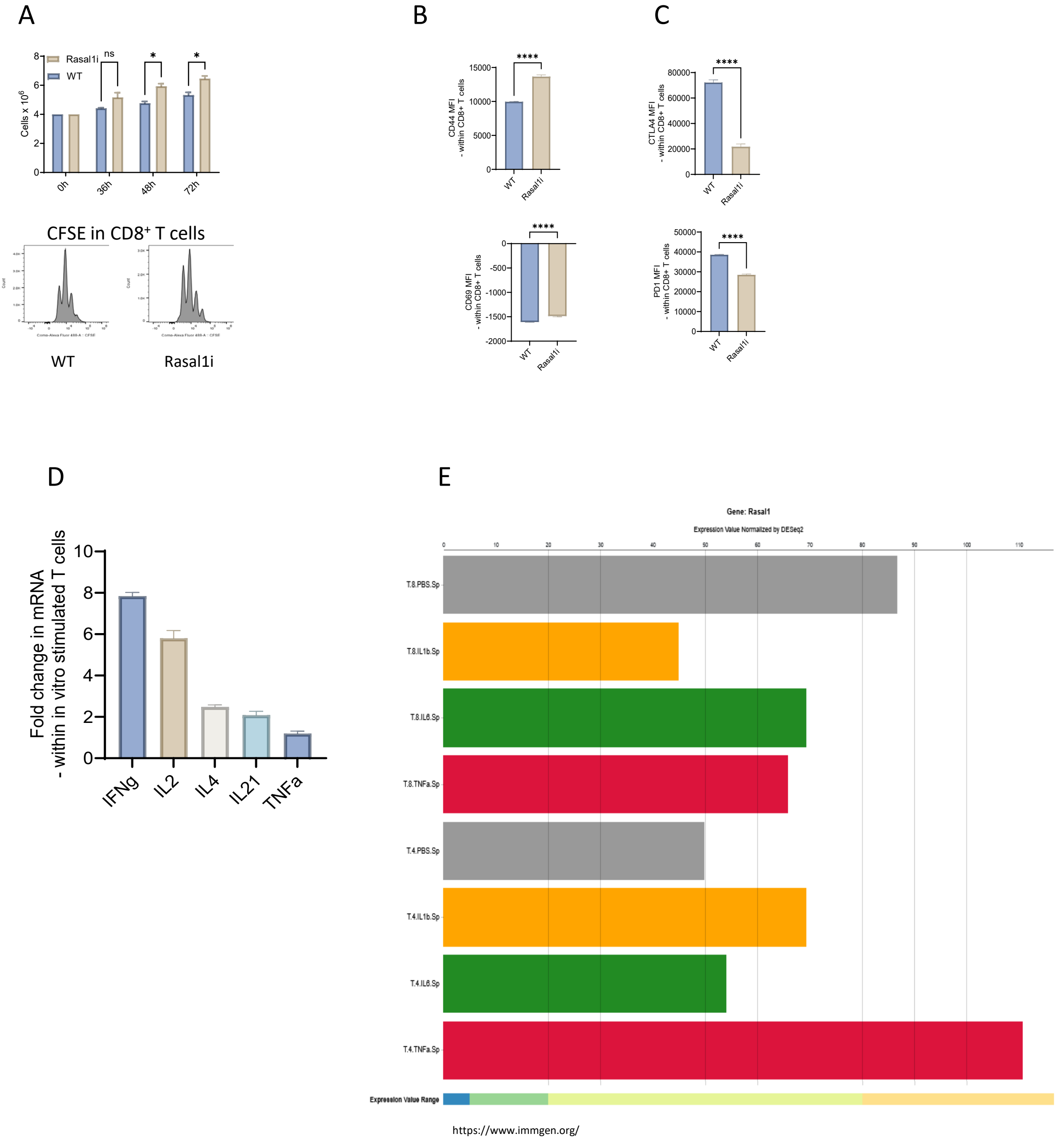

### Supplemental Figure 2

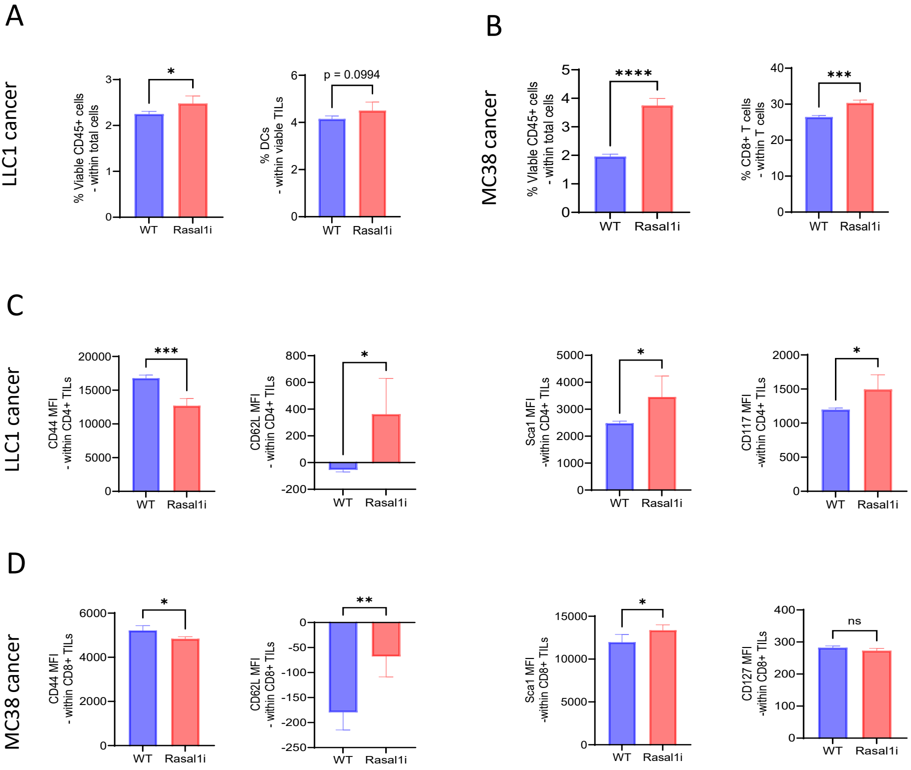
